# Identification of Significant Computational Building Blocks through Comprehensive Investigation of NGS Secondary Analysis Methods

**DOI:** 10.1101/301903

**Authors:** Md Vasimuddin, Sanchit Misra, Srinivas Aluru

## Abstract

Rapid advances in next-generation sequencing technologies are improving the throughput and cost of sequencing at a rate significantly faster than the Moore’s law. This necessitates equivalent rate of acceleration of NGS secondary analysis that assembles reads into full genomes and identifies variants between genomes. Conventional improvement in hardware can at best help accelerate this according to the Moore’s law. Moreover, a majority of the software tools used for secondary analysis do not use the hardware efficiently. Therefore, we need hardware that is designed taking into account the computational requirements of secondary analysis, along with software tools that use it efficiently. Here, we take the first step towards these goals by identifying the computational requirements of secondary analysis. We surveyed dozens of software tools from all the three major problems in secondary analysis – sequence mapping, *De novo* assembly, and variant calling – to select seven popular tools and a workflow for an in-depth analysis. We performed runtime profiling of the tools using multiple real datasets to find that the majority of the runtime is dominated by just four building blocks – Smith-Waterman alignment, FM-index based sequence search, Debruijn graph construction and traversal, and pairwise hidden markov model algorithm – covering 80.5%-98.2%, 63.9%-99.4% and 72%-93% of the runtime, respectively, for sequence mapping, *De novo* assembly, and variant calling. The key outcome of this result is that by just targeting software and hardware optimizations to these building blocks, major performance improvements for NGS secondary analysis can be achieved.

## 1 Introduction

The invention and rapid advancements of Next Generation Sequencing (NGS) technology has spurred advancements in fields such as genetic testing, DNA based disease diagnosis and treatment, gene editing, etc., with applications to human health (e.g. personalized medicine), agriculture, archeology and forensics. This revolution is best exemplified by Illumina sequencers. A single Illumina Hiseq X 10 system, released in 2014, can sequence nearly 18000 human genomes per year, at the low cost of less than $1000 per genome, sequencing short DNA fragments (called reads) of length 150 bp at the rate of nearly 1.6 quadrillion bp per year [29]. Moreover, the recently released Illumina NovaSeq 6000 system [30] is nearly 3.75X faster than Illumina HiSeq X system. High throughput and low cost have led to the widespread adoption of these sequencers with large sequencing centers employing dozens of them. This has ushered in the era of million human genomes, with several countries and public and private organizations pursuing sequencing genomes of a million or more humans [1,3,13,16,21,22,32,37,63] to enable population level studies. It is estimated that genomes of up to 2 billion humans could be sequenced by 2025 [62].

**NGS** based genomic studies are performed in three stages as follows. *Primary analysis* is performed by sequencers. Given a biological sample, typically, multiple copies of the genome sequence contained in it are extracted and then decomposed into smaller fragments. A sequencer reads the sequence of nucleotides in the fragments and generates signals based on what it reads. These signals are interpreted to derive reads as sequences of bases over the nucleotide alphabet {A,C,G,T}, and the corresponding quality score for each base. Thus, the reads correspond to substrings of the genome. The sequencer outputs these reads and quality scores and they are represented across all sequencers using the same uniform representation, typically a **FASTQ** file [14]. In *secondary analysis*, one of the fundamental tasks is to construct the complete **DNA** sequence from the reads. This is typically done by mapping the reads to one or more reference genomes, or assembling them *de novo* based on read overlaps in the absence of a suitable reference. Another crucial task is to identify variants with respect to a reference or among the samples. After getting the variants, *tertiary analysis* works on understanding the implications of those variants on the study of interest.

The sequencers are getting faster and cheaper at an exponential rate much faster than the Moore’s law. Given the rapid pace of sequencers and the ambitious goals like sequencing millions of genomes, commensurate speeds are required for **NGS** secondary and tertiary analysis. Conventional architectural improvements can at best help accelerate these according to the Moore’s law. Therefore, we will require efficient use of the current architectures, and new architectures that are tailored to achieve high performance for **NGS** secondary and tertiary analysis.

While tertiary analysis is still an emerging field, significant developments have already taken place in secondary analysis. Therefore, in this paper, we focus on **NGS** secondary analysis. There are hundreds of software tools available for **NGS** secondary analysis. Even if we restrict to only the most widely used tools, there are still at least a dozen of them. Moreover, given the dynamic nature of the field, the most widely used tools constantly get modified or replaced by newer tools. This makes it impractical to accelerate the tools or design architectures for them. However, while the tools keep changing, the underlying key computations seem restricted to a small set of building blocks. Accelerating these building blocks can have a significant impact on the performance of **NGS** secondary analysis. Thus, we focus this study on the identification of computational building blocks of **NGS** secondary analysis. Our work can inform any future efforts to accelerate **NGS** secondary analysis through improvements in algorithms, software, and hardware.

To identify the building blocks, we performed a rigorous survey of the secondary analysis methods to select tools and techniques for our study. We used the following criteria for the selection – the tools should be high-quality, well-maintained, and widely used. We downloaded the latest source of the tools, studied the source codes to understand them in full detail to identify various common building blocks across tools, and hand instrumented them with runtime profiling instructions. We followed the instructions given in the documentation of the tools to build, install, and run the tools. The evaluation was carried out using real datasets. Overall, we studied seven tools and a workflow from the three primary areas of *De novo* assembly, sequence mapping, and variant calling in secondary analysis. The evaluation of the runtimes revealed following four primary building blocks: Smith-Waterman sequence alignment, **FM**-index based sequence search, Pairwise hidden markov model algorithm for sequence alignment likelihood calculations (PairHMM), and de Bruijn graph for *De novo* assembly. Together, these building blocks cover 63.9%-99.4% of time for *De novo* assembly, 80.5%-98.2% of time for sequence mapping and 72%-93% of time for variant calling. The implication of this result is that by just tailoring software and hardware advancements to target these building blocks, major performance improvements in **NGS** secondary analysis can be obtained. To the best of our knowledge, there has not been a comprehensive effort to study a wide range of software tools for **NGS** secondary analysis that tries to carve out similar building blocks across them by studying the source code of each of them in detail, and formally establish the building blocks with profiles generated using real datasets.

The rest of the paper is organized as follows. In section 3, we discuss components of secondary analysis and outline our strategies for selecting workflows and software tools. The following sections contain indepth analysis and profiling results for sequence mapping (Section 4), *denovo* genome assembly (Section 5), and variant calling (Section 6). Additional discussion and conclusions are presented in Sections 7 and 8, respectively.

## 2 Experiment Setup

We use the following experimental setup for evaluation of all the tools and workflows studied in this paper.

All the single node experiments along with multi-node experiments for ABySS (section 5) were carried out on a cluster with 16 nodes with each node comprising of a dual-socket Intel^®^ Xeon^®^ processor with 18-cores per socket (HE5-2699). Each compute node is equipped with 128GB of memory, running CentOS Linux version 7.2. The compute nodes are interconnected using Infiniband FDR interconnect. Multi-node experiments for HipMer (section 5) were performed on NERSC’s Cori supercomputer. Each compute node is equipped with a dual-socket Intel^®^ Xeon^®^ processor with 16-cores per socket (HE5-2698 v3) along with 128GB of memory. The compute nodes are interconnected by Cray Aries interconnect with Dragonfly topology.

## 3 Secondary Analysis Methods

**NGS** secondary analysis takes raw reads generated from a sample using primary analysis as input and outputs the corresponding genome and variants compared to other genome(s). The first step is to reconstruct the genome by stitching together the reads by either mapping them to a reference genome, called sequence mapping, or assembling them *de novo* by leveraging the coverage depth and overlap information among the reads.

Given genomic datasets, a crucial task in sequence analysis is finding the differences among the genomes. Variant calling (**VC**) aims at precisely finding the genomic locations exhibiting variations such as single nucleotide variants, short indels, and large structural variants. Variants in a sequenced genome with respect to a reference genome can be found by mapping the reads directly to the reference. Another alternative is to first reconstruct the sequenced genome directly from the reads through *de novo* genome assembly. Sequence mapping based assemblies are biased towards the reference genome. However, generating high-quality genomes using *de novo* assembly is both difficult and extremely time consuming. Consequently, almost all **VC** workflows use sequence mapping as the preprocessing step. However, this can change in the future or the use of *de novo* assembly may be more appropriate in certain contexts. Hence, we consider software for both approaches in this work.

**VC** workflows are sequences of steps, each performed by a software tool, that need to be executed to go from reads to variants. The choice of tools used in such workflows depends on the type of application. The high impact of, and the challenges posed by, **VC** has prompted the development of numerous **VC** tools and workflows. Among the various **VC** calling workflows such as Samtools [41], **GATK** (Genome Analysis ToolKit) [18] best practices workflows, Platypus [57], and DeepVariant [54], **GATK** best practices workflows developed at Broad Institute are by far the most popular and actively-maintained workflows. They are hosted on widely used platforms including Google cloud, Microsoft Azure, Amazon web services, Ali cloud, etc., and are used by researchers all over the world. **GATK** best practices workflows have been widely adopted by the community due to its ability to handle large-scale **VC** studies and to identify high quality variants. Recently proposed convolution neural network based DeepVariant workflow demonstrated impressive results; however, further studies are required to convincingly establish its practical applicability. Therefore, in this work, we focus on **GATK** workflows; we begin with a study and runtime profiling of **GATK** workflows to highlight the compute-dominant stages.

### 3.1 GATK Best Practices Workflows

**GATK** encompasses a suite of software tools targeted towards studying different kinds of variants such as SNP/indels/copy number variations (CNV) of *germline* or *somatic* type. In addition to the software, **GATK** is augmented with a wealth of literature and best practice guidelines [8] for using the workflow of interest. Among the matured workflows in **GATK**, workflows utilizing HaplotypeCaller tool are suitable for detecting *germline* SNP/indel, while workflows utilizing MuTect2 tool are suitable for detecting *somatic* SNP/indel. *MuTect2* tool has majority of its operations similar to *HaplotypeCaller* tool; with *MuTect2* borrowing the assembly based engine of *HaplotypeCaller* to original *MuTect* [12]. Considering the similarity between the two tools, and the maturity and high-quality variant detection capability [28,51] of HaplotypeCaller, in this work we focus on the **GATK**-HaplotypeCaller workflow.

Next, we describe different stages of the workflow and the tools recommended for executing them (Figure 1). Given a reference genome and read set, the workflow executes the following steps:

- *Sequence Mapping*. For each read, sequence mapping outputs a set of locations in the reference genome where the read aligns to the reference genome sequence while allowing a few mismatches and gaps in the alignment. **GATK** best practices recommends available prominent sequence mapping tools such as **BWA-MEM** [38].
- *Sorting*. With genomic locations of the reads known, reads are sorted using genome-coordinates, to cluster together reads mapped to the same region in the reference genome. This also helps in identifying duplicate reads. **GATK** best practices recommends Picard’s [53] *SortSam* tool for this step.
- *Mark Duplicate Reads*. Duplicates are reads that are likely to have originated from duplicates of the same original **DNA** fragments. Each read provides independent evidence towards identifying the variations; however, duplicate reads offer no additional information about the variations but can artificially skew the support towards a particular variant and also require more time in processing. Thus, duplicate copies are marked and excluded from further processing. **GATK** recommends Picard’s *MarkDuplicates* tool for this step.
- *Base Recalibration*. Base quality scores are extensively incorporated during variant discovery. However, the quality scores reported by the sequencers contain technical errors which can be estimated and fixed. This workflow first models the error patterns in the data using its *Base Recalibrator* tool.
- *Recalibrate Base Scores*. **GATK** provides *PrintReads* tool which adjusts the base quality score, based on the model learned in the previous step. The best practices categorizes all the processing until this step as data *pre-processing*. The pre-processed data is then ready for variant detection.
- *Variant Discovery*. Given pre-processed data, variant discovery identifies genomic locations exhibiting variations. **GATK** provides *HaplotypeCaller* (**HC**) [55] tool for *germline* **VC**. The variants reported by the *variant discovery* step are subjected to further post-processing which is categorized under tertiary analysis.

**Figure 1.**
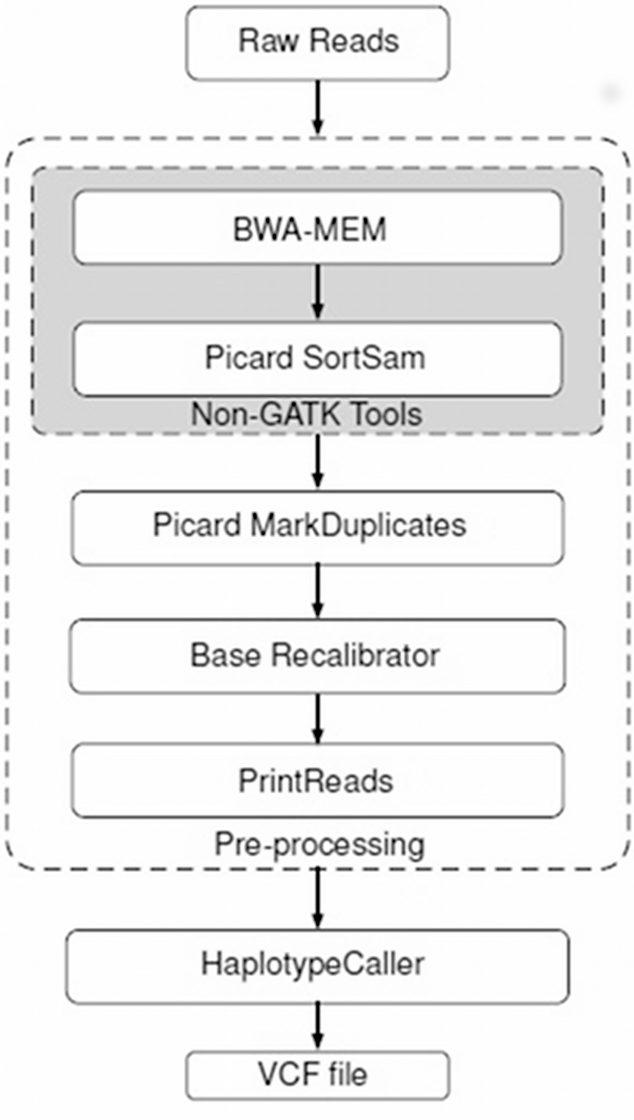
**GATK**-3.8 best practices workflow for *germline* **VC**.

Different tools in the workflow support different types of parallelism. **BWA-MEM** and *PrintReads* support thread-level parallelism, while *BaseRecalibrator* and *HaplotypeCaller* support parallelism using *scatter-gather* method. The *scatter-gather* method works as follows: In the scattering phase, data is partitioned into multiple smaller parts. These parts are worked on using process-level parallelism in which individual process, with its memory space, is spawned for each part of the partitioned input data. At the end of the execution results from all these processes are gathered to report the combined results.

We used **GATK** version 3.8, the most recent version available at the time of this study that has full documentation available, to perform runtime profiling of the workflow. We used human genome version 38 (NCBI) as the reference genome, and low coverage (GBR population, identifier HG00119) read sets acquired from NCBI’s Sequence Read Archive (SRA, accession no. SRX020470 and SRX020450). The read sets are sequenced by the Broad Institute using Illumina Genome Analyzer-II [6]. The datasets contain seven paired-end read sets with each read set containing on average 34 million reads, collectively containing 250 million reads.

The end-to-end sequential execution of **GATK**’s workflow on given input datasets took 33.31 hours on a single core (Table 1). The runtime profile of the workflow revealed that *Sequence Mapping* and *Variant Discovery* are dominant stages, consuming 52.5% and 27.3% receptively, of the total runtime. Similar observations are also witnessed by a study [56] on multiple human genome-scale datasets. For multi-core parallel runs, we followed the **GATK** recommendations and, for each tool, we experimented with different number of cores. We present the best performing multi-core result for each tool in the table. **BWA-MEM** is able to scale well up to all the 36 cores of single node. The Picard tools SortSam and MarkDuplicates do not have parallel implementations and thus can utilize only a single core. However, various parallel implementations are available for sorting, while *MarkDuplicates* operation offers ample parallelism to accelerate it, which would lead to significant drop in the time consumption of these tools. Even though *PrintReads* consumes significant enough time to be considered important for acceleration, in the recent release of **GATK** (version 4.0), it has been replaced with *ApplyBQSR* that is significantly faster making it relatively unimportant for this study. Thus, with *BaseRecalibrator* consuming relatively small time, we are left with *Sequence Mapping* and *Variant Discovery* as the compute dominant stages. Therefore, we narrow our search for building blocks in **VC** to *Sequence Mapping* tools (Section 4) and *Variant Discovery* tools (Section 6), along with *De novo* assembly (Section 5).

**Table 1.** Time spent in different tools of end-to-end execution of **GATK**’s *germline*-**VC** workflow on human genome dataset. The numbers in the brackets in single-node runs specify the maximum number of cores utilized by the tools.

| Tools | Single-core |  | Single-node |  |
| --- | --- | --- | --- | --- |
|  | Time (h) | Time (%) | Time (h) | Time (%) |
| BWA-MEM | 17.50 | 52.54 | 0.70 (36) | 10.82 |
| Picard SortSam | 0.68 | 2.04 | 0.68 (01) | 10.51 |
| Picard MarkDuplicates | 0.65 | 1.95 | 0.65 (01) | 10.05 |
| BaseRecalibrator | 2.00 | 6.00 | 0.24 (28) | 3.71 |
| PrintReads | 3.40 | 10.20 | 1.55 (08) | 23.96 |
| HaplotypeCaller | 9.08 | 27.26 | 2.65 (07) | 40.96 |
| Overall runtime | 33.31 |  | 6.47 |  |

## 4 Sequence Mapping

Given a reference genome and a set of reads, sequence mapping or alignment finds the probable locations for each of the reads in the reference genome. Many workflows use sequence mapping as the first step in VC.

Most modern sequence mapping tools use the *seed-and-extend* strategy to map a read to a reference sequence. In the seeding stage, they find regions in the reference sequence that closely match subsequences from the read, called seeds. In the extend stage, these regions are evaluated more closely to verify if they are a good match of the entire read. A majority of tools use dynamic programming (**DP**) based algorithms for extension. On the other hand, data structures play a central role in seeding, primarily used for indexing either the reference genome, or the read sequences, or both.

Many sequence aligners have been proposed over the years, utilizing different data structures. Hash-based aligners [27,31,42,43,46,58,61] are used to hash *k*-mers either in the reference or in the reads. Seeding stage uses the hash table to find regions in the reference that have matches for *k*-mers from the read or their minor modifications. Suffix trie is another important data structure used for indexing the sequences [4,26,33,50]. A suffix trie stores all the suffixes of a sequence *S*, such that each edge is labeled by a character from *S*. Any path from the root to a leaf in the suffix trie corresponds to a suffix of *S* and from root to an internal node corresponds to a substring of *S*. A seed is searched in a trie by traversing it from the root using the characters of the seed. Methods using hash-table or trie are expensive in terms of memory usage.

Most prominent mapping software use either the space-efficient Burrows Wheeler Transform (**BWT**) [9], or the BWT-based **FM**-index data structure proposed by Ferragina and Manzini [20] for seeding. For a conceptual understanding of the **BWT** of a string *S* of length *n* – 1, consider appending the lexicographically smallest character $ to *S*. Consider the *n* × *n* matrix obtained by listing the *n* rotations of the appended string as its rows. The **BWT** matrix is the resulting matrix when these rows are lexicographically sorted. The last column of the **BWT** matrix represents the **BWT. BWT** has an interesting property termed *last-first*, or **LF** mapping, which preserves the order of instances of character *X* between the last and the first column of the matrix. The **LF** property led Ferragina and Manzini to create the **FM**-index and derive an exact string matching method based on it. The **FM**-index maintains additional auxiliary arrays including suffix array (**SA**). **SA** stores the reference locations for each row (i.e.suffix) of the matrix. Query sequences are passed through the **FM**-index in reverse order (called *backward search*), which is equivalent to top-down traversal on a prefix trie. Memory footprint of **FM**-index is very small (less than a few GB), which makes it a highly practical and preferred choice over the other data structures. Many sequence mapping tools [34–36,38–40,44] based on **BWT** have been proposed over the years, of which **BWA-MEM** [38] (recommended by **GATK** best practices workflow) and Bowtie2 [35] are by far the most popular due their speed and accuracy. Hence, we selected Bowtie2 and **BWA-MEM** for our study.

### 4.1 Chosen Sequence Mapping Tools

Bowtie2 and **BWA-MEM** also use the popular *seed-and-extend* strategy. Both the tools use bi-directional **FM**-index of the reference sequence for seeding, and dynamic programming (**DP**) based alignment methods for extending the seeds. The tools differ in their approaches for seeding and extension.

For a given read, Bowtie2 extracts seeds (substrings of reads) of a particular length at a regular interval from the read and its reverse complement. It offers the options of exact matching as well as inexact matching (with 1-mismatch) of seeds in the references sequence.

The output of the *seeding* stage is a list of regions in the reference genome where at least one seed matches, called candidate regions. For each candidate region, the *extension* phase verifies whether the region is a good match of the read or not. During *extension*, Bowtie2 uses a **DP** technique based on the Smith-Waterman algorithm for sequence alignment. The **DP** technique computes a two-dimensional matrix between the reference and query sequences. At the end of the matrix computations, it reports the high scoring alignment. For paired-end reads, Bowtie2 follows the *seed-and-extend* steps for each end individually. Once the first end of the paired-end read is aligned, based on the insert distance, Bowtie2 computes the reference window which contains the probable location for the second end. The sequence alignment is performed for the second end in the identified window.

**BWA-MEM** performs seeding by finding *super maximal exact matches* (**SMEMs**) between the read and the reference sequence using **FM**-index. For a given position in the read, the **SMEM** corresponding to that position is the longest exact match through that position. For paired-end reads, **BWA-MEM** sorts all the **SMEMs** according to the genome coordinates to identify the **SMEMs** corresponding to paired-end reads. In the extension phase, **BWA-MEM** uses **DP** based banded Smith-Waterman algorithm. In banded Smith-Waterman, during the matrix computations, only the cells falling within certain band size around the diagonal are computed. **BWA-MEM** also applies the following additional heuristics to **DP** computations. (a) During matrix computations, if the score falls significantly below the best score, the execution halts; (b) if the difference between the best local alignment score and the global alignment score is below a certain threshold, then the best local score is discarded in favor of the global score. These heuristics reduce computation and provide control over reference bias.

### 4.2 Results

For both Bowtie2 and **BWA-MEM**, we downloaded the latest available source codes (Bowtie2-2.3.2 and **BWA-MEM**-0.7.15) for our evaluation, studied them in full detail to identify boundaries of similar building blocks, hand instrumented them with runtime profiling instructions, and followed the recommendations given in the corresponding *readme* files to install, compile, and run them. We also used recommended default parameter settings. We used two different datasets for evaluating the software. The first consists of full human reads sets (identifier HG00119) that we also used for profiling **GATK**’s best practices workflow. The second is a low coverage (CEU population, identifier NA12878) single-end read set acquired from NCBI’s Sequence Read Archive (SRA, accession no. SRX206890). The read set contains 1.4 billion reads; however, due to high compute demands of inexact matching in Bowtie2, we uniformly sampled 30 million reads from this dataset to use for this study.

Figures 2 & 3 show the percentage of the overall runtime consumed by each block in Bowtie2 and **BWA-MEM**, respectively. Seed matching using **BWT**, *SA2Ref*, and seed extension using Smith-Waterman (**SWA**) appear to be the runtime dominant blocks for both software tools. Given a seed, once the row range in **BWT** matrix is computed, *SA2Ref* looks up the **SA** entries in the **FM**-index corresponding to the range to find the corresponding reference genome locations. Blocks that are not profiled are denoted as *Misc*. Bowtie2 also allows one mismatch for seed search spending a lot more time in **BWT** in that case due to a significantly larger search space.

**Figure 2.**
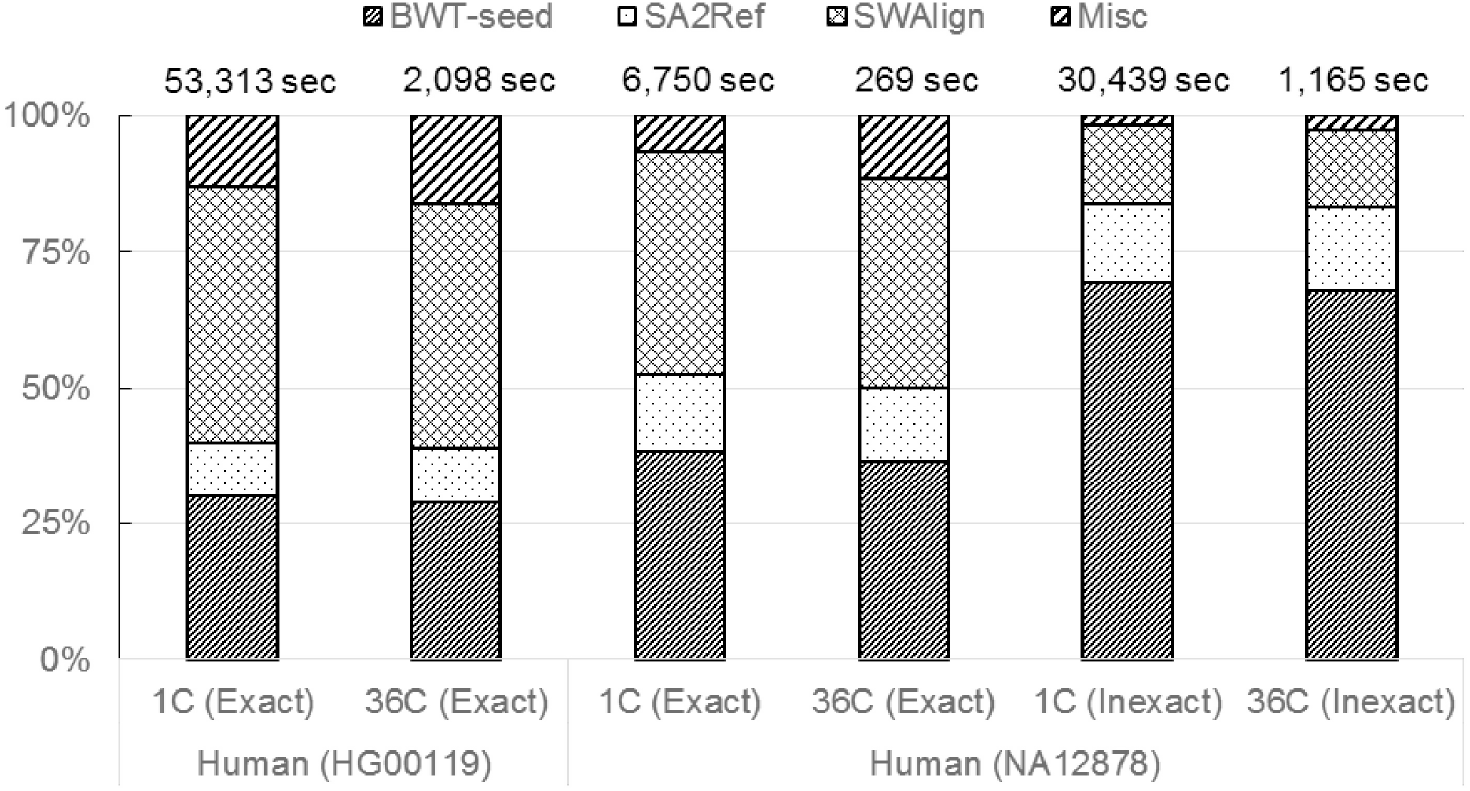
Percentage of the overall runtime consumed by different blocks of Bowtie2 for single as well as 36 cores (C). Exact and Inexact refer to exact seed matching and seed matching with 1 mismatch allowed, respectively.

**Figure 3.**
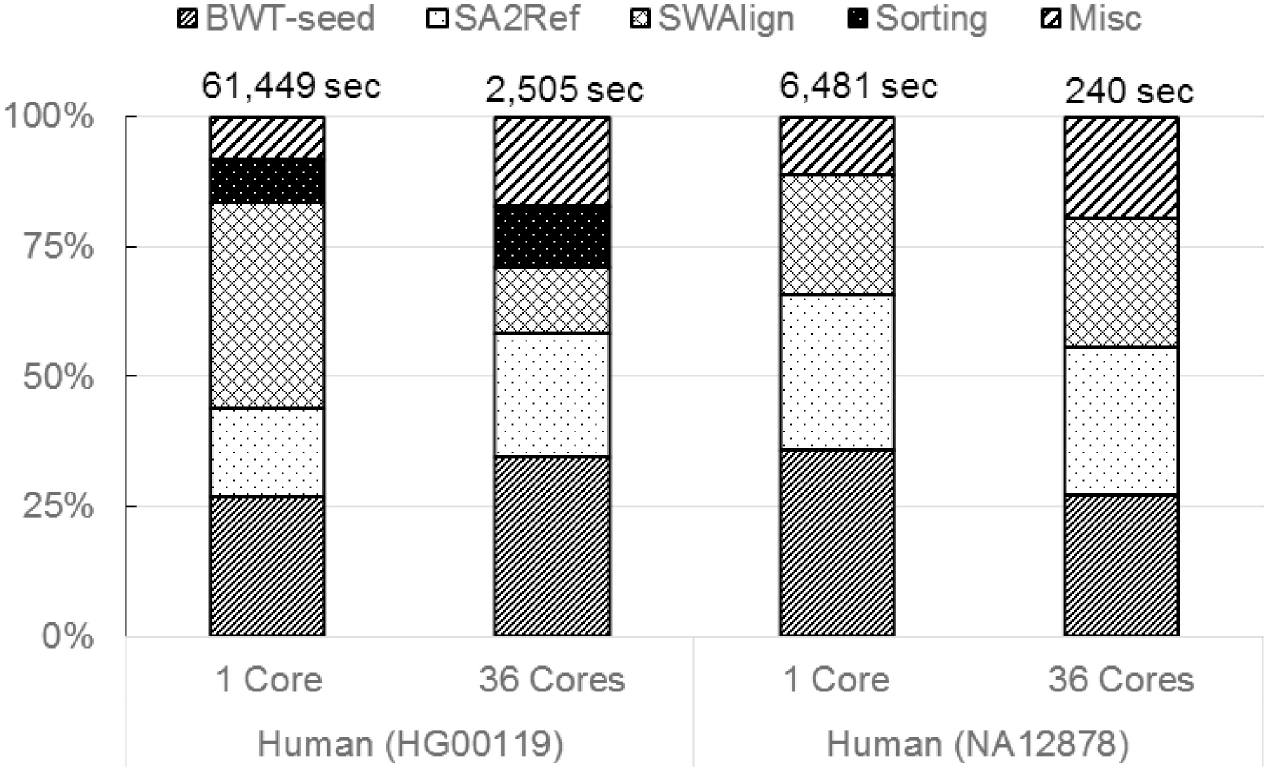
Percentage of the overall runtime consumed by different blocks of BWA-MEM for single as well as 36 cores.

*SA2Ref* and **BWT** together constitute the building block **FM**-index based sequence search that uses the **FM**-index of a sequence to find the positions of all the occurrences of another sequence in it. Our runtime profiling shows that **FM**-index based sequence search and **SWA** constitute the most significant blocks in run-time consumption for both the sequence mapping tools for single as well as multiple threads covering 80.5% – 98.2% of the total time. Thus, we categorize **FM**-index based sequence search and **SWA** as key building blocks.

## 5 De novo Assembly

Constructing genomes directly from raw reads is a fundamental step in studying the genome of an organism, and is the only viable path in the absence of a suitable reference sequence. Factors such as small read lengths, large number of reads, sequencing errors, and genomic repeats make *de novo* genome assembly extremely challenging. Pevzner et al. [52] successfully applied de Bruijn graphs for genome assembly, which has since been used by a majority of short read genome assemblers.

Numerous assemblers have been proposed over the years, leading to the establishment of competitions such as Assemblathon [2] and GAGE [48,59] to benchmark their accuracy and performance. These competitions extensively evaluate submitted assembly workflows over a range of metrics using benchmark datasets. Recently held Assemblathon-II competition concluded that majority of the assemblers perform better than others only on particular subsets of metrics and datasets. Thus, in a scenario where choice of assemblers heavily relies on the type of study, we picked a few assemblers that are mature and favored among the assembly workflows benchmarked in such competitions. We observed that among the genome assembly workflows submitted to Assemblathon-II and GAGE-A/ B, ABySS [60], SOAPDenovo [45,47], SPAdes [5], Meraculous (HipMer) [11,24], AllPaths [10], Ray [7], and Velvet [64] consistently performed better over a range of metrics and were highly preferred. From these assemblers, we selected a blend of distributed and shared memory assemblers which are popular and widely used. Since it is difficult and unnecessary to study all the high performance assemblers, we restricted the number of assembler for out study to the following four: ABySS, SOAPDenovo, SPAdes, and Meraculous (HipMer). Given a reads set, a majority of the new generation assemblers use the following assembly framework.

1. *Graph Construction*. De Bruijn graph is a directed (or bidirected when both **DNA** strands are directly modeled) graph, with *k*-mers as nodes and edges connecting *k*-mers that share (*k*-1) length suffix-prefix overlap. A de Bruijn graph can be stored in a hash-table. *k*-mers are used as keys and the counts of each extension (i.e.A,C,T,G) of *k*-mers are stored as values in the hash table.
2. *Graph Cleaning*. Errors in reads manifest as various structural artifacts such as bubbles, tips, tiny repeats, etc., in the constructed graph. As a result, such artifacts need to be located and removed before extraction of contigs. Graph cleaning is accomplished by traversing the graph.
3. *Contig Extraction*. The graph is traversed along unambiguous paths, generating initial contigs.
4. *Read Alignment*. Information contained in paired-end reads can be utilized to extend the contigs. As a first step, paired-end reads are aligned to their corresponding contigs. The alignment helps identify contig orientation and ordering.
5. *Scaffolding*. Contigs corresponding to paired-end reads are identified. Link between the contigs is created if a certain number of read pairs support it.
6. *Gap Closer*. Gaps between contigs in a scaffold primarily contain repetitive regions. Paired-end information can be used to fill the gaps. Paired-end reads having one read mapped to a contig and the other falling in a gap are retrieved. The reads corresponding to the gaps are then *de novo* assembled to fill the gaps. In the presence of multiple insert libraries for paired-end reads, the libraries are utilized iteratively from smaller to larger insert sizes. This step is performed iteratively.

In the above framework, the first three steps can be performed using reads without pairing information, while paired-end reads are required to expand the initial contigs. Three of the four assembly programs we have selected for our study follow the above framework, while SPAdes assembler employs *paired* de Bruijn graphs.

### 5.1 Assembly Methods and Software

We briefly describe the methodologies behind the four assembly software selected for the study.

#### ABySS

*De novo* assembly is memory intensive, and for assembling mammalian sized genomes large read sets must be analyzed. To ease the memory limitations, Simpson **et al**. [60] proposed ABySS, a distributed-memory based parallel assembler where one core runs one process. A key aspect of ABySS is the distributed-memory construction of the de Bruijn graph, constructed as follows. Given the reads, the extracted *k*-mers are converted to numerical values by assigning (0,1,2,3) to (A,C,G,T) bases, and treating them as base 4 numbers. The resulting *k*-mer values are hashed, and distributed to cores based on their values modulo *P* (total number of cores in the distributed system). For each *k*-mer, 8 bits are used to store its adjacency information. Each bit represents the presence or absence of each possible extension (**i.e.**an edge) on either side of the *k*-mer. After graph construction, contigs are extracted by traversing the graph in parallel. ABySS uses sequential computations to execute the rest of the steps in the assembly framework.

#### SOAPDenovo

SOAPDenovo [47] is a shared-memory genome assembler that performed very well in Assemblathon-I competition ranking overall second on evaluation in eight categories. SOAPDenovo2 [47] further improves upon SOAPDenovo on various fronts, as follows: (a) Memory consumption of SOAPDenovo, a bottleneck, is mainly contributed by the graph construction stage. SOAPDenvo2 constructs a sparse de Bruijn graph instead, which reduces memory footprint by grouping linear chains of *k*-mers, thus avoiding the need to store each *k*-mer separately. (b) The choice of *k*-mer size impacts de Bruijn graph construction. For assembly, smaller *k*-mer sizes offer advantage in low coverage regions, while larger *k*-mer sizes are useful to handle repetitive sequences. SOAPDenovo2 avails the benefits of multiple *k*-mer sizes, by iteratively building de Bruijn graph using different *k*-mer sizes. (c) To improve quality of the scaffolds, SOAPDenovo2 performs additional processing to ease the effects of heterozygosity, chimeric scaffolds, and false contig links during scaffolding. (d) SOAPDenovo2 improves the accuracy of the *Gap Closer* step in highly repetitive regions. At each iteration in *Gap Closer*, in addition to the reads that map to the gaps in the current iteration, SOAPDenovo2 also utilizes mapped reads from the previous iterations to fill the gaps.

#### SPAdes

Bankevich **et al**. proposed the SPAdes genome assembler for shared-memory parallel systems, designed to overcome the challenges posed by both single-cell as well as multi-cell genomes. SPAdes deviates from common assembly framework which utilizes paired-end reads (referred to as *bireads*) only after contig extraction. After constructing the de Bruijn graph from reads, SPAdes applies the concept of *paired* de Bruijn graph (**PDBG**) by Medvedev **et al**. [49], utilizing biread information to generate the final contigs. While distances between the reads in bireads are only known approximately, the construction of **PDBG** requires knowledge of exact distances. SPAdes addresses this issue by estimating the distances.

SPAdes execution has the following four major stages. (a) SPAdes iteratively constructs the de Bruijn graph by utilizing multiple *k*-mer sizes; the constructed graph is referred to as *multisized* de Bruijn graph. Given multiple *k*-mer sizes, the *multisized* de Bruijn graph is constructed as follows. For a given *k*-mer size, a standard de Bruijn graph is constructed, and all the paths having vertices with in- and out-degree as 1 (referred as *h-edges*) are extracted from it. The extracted *h-edges* are thus included as reads for the next iteration of graph construction. Using *multisized* de Bruijn graphs, SPAdes simultaneously exploits the advantages of both small and large *k*-mer sizes by increasing value of *k*-mer at each iteration. (b) A major impediment to generate the PDBG is finding the exact distances within bireads. SPAdes uses a computational method based on the Fast Fourier Transform to estimate these distances within the bireads (or *k*-bimers, pair of *k*-mers). The *k*-bimers are then adjusted according to the estimated distance. (c) The **PDBG** is then constructed by using the estimated distances and adjusted *k*-bimers. For more details, the reader is referred to [49]. (d) Contigs are then extracted by traversing the **PDBG**.

#### HipMer

HipMer [24] is an extreme-scale distributed-memory parallel implementation of the Meraculous assembler [11]. Meraculous follows steps of the outlined assembly framework with one distinction: it counts the frequency of occurrence of individual *k*-mers and discards low frequency *k*-mers by classifying them as erroneous, instead of identifying and removing errors through topological features of the graph. HipMer employs novel techniques and parallelizes each assembly step to scale to thousands of processors. High communication overhead degrades throughput during distributed graph traversal. HipMer overcomes the challenge by applying a communication avoidance algorithm. The algorithm derives and makes use of a partitioning function. The partition function allocates *k*-mers to processors such that *k*-mers belonging to the same contig are likely allocated to the same processor. HipMer generates the partitioning function by exploiting the fact that genomes of different individuals of the same organism are highly similar.

After contig extraction, Hipmer maps the reads to contigs using the parallel sequence aligner, mer-Aligner [25]. The contigs are then connected to form scaffolds, using the aligned paired-end reads. For gap closing, HipMer employs multiple techniques depending on the complexity of the gaps. Gap closing is parallelized by equally dividing the gaps among the processors. Each processor applies the following methods in succession to fill the gaps. *Spanning*, which finds out the reads which overlap with a contig tail at one end of the gap and another contig’s head at the other end; *mini-assembly*, which re-assembles the reads aligned to the gap regions and traverses the graph to find the fillers; and *Patching*, for cases when graph traversal from both the ends of a gap fails, then HipMer tries to patch the two traversals by finding the overlap between them.

### 5.2 Results

We downloaded the latest available source codes of all the assemblers – ABySS-1.9.0, SOAPDenovo-2.0, SPAdes-3.10.1, and HipMer-0.9.4.1, performed an in-depth study on them to identify boundaries of similar blocks and hand instrumented them for runtime profiling. We followed the instructions from *readme* files to install, compile, and run the programs. Each software tool is run using standard benchmark datasets from GAGE [59] (Table 2), using default parameters.

**Table 2.** Benchmark short-reads datasets for **VC** *De novo* assembly (M-Millions). Chr14 refers to the chromosome 14 of the human genome.

| Datasets | Library | No. of reads | Avg. read len. | Insert len. |
| --- | --- | --- | --- | --- |
| Ecoli | Frag | 28M | 101bp | 215bp |
| Chr14 | Frag | 36M | 101bp | 155bp |
|  | Short | 22M | 101bp | 2283-2803bp |
|  | Long | 2M | 76-101bp | 35,295-35,318bp |
| Bumble Bee | Frag | 1303M | 124bp | 400bp |

For each assembler, we used at least two datasets and conducted experiments using small as well as a large number of cores, to study changes in the runtime of the blocks with change in scale of data or the hardware used. For single node experiments, we used the smaller datasets *E. coli* and Human chromosome-14 (Chr14). For Abyss and Hipmer that have support for distributed memory systems, we used the larger Bumblee Bee dataset to perform experiments using higher number of nodes. Moreover, as the choice of *k*-mer size affects the runtime of an assembler, we experimented with different *k*-mer sizes for all the assemblers except SPAdes. SPAdes performs automatic selection of *k*-mer size based on the read length. While we noticed different runtimes for different *k*-mer sizes, the proportion of the total runtime for the individual blocks remained approximately the same. Therefore, we only report the performance for one *k*-mer size.

Figures 4-7 show the percentage of overall runtime consumed by each block for the four assembly software. De Bruijn graph construction is used by all the four assemblers and consumes a major portion of the overall runtime for each. Except for SPAdes, all the other assemblers follow the framework described in section 5.1; and for them, *Sequence alignment* and *gap closer* blocks are the other two primary consumers of the overall runtime. *Gap closer* iteratively performs assembly over the gap regions. Thus, we do not classify it as a separate building block. For SPAdes, *k-mer Adjustment* forms the second big block; however, runtime consumed by it remains relatively small. As before. the runtime spent in portions of the code that are not profiled is labeled as *Misc*. It is clear that the runtime of major blocks remains dominant with the change in the number of cores.

**Figure 4.**
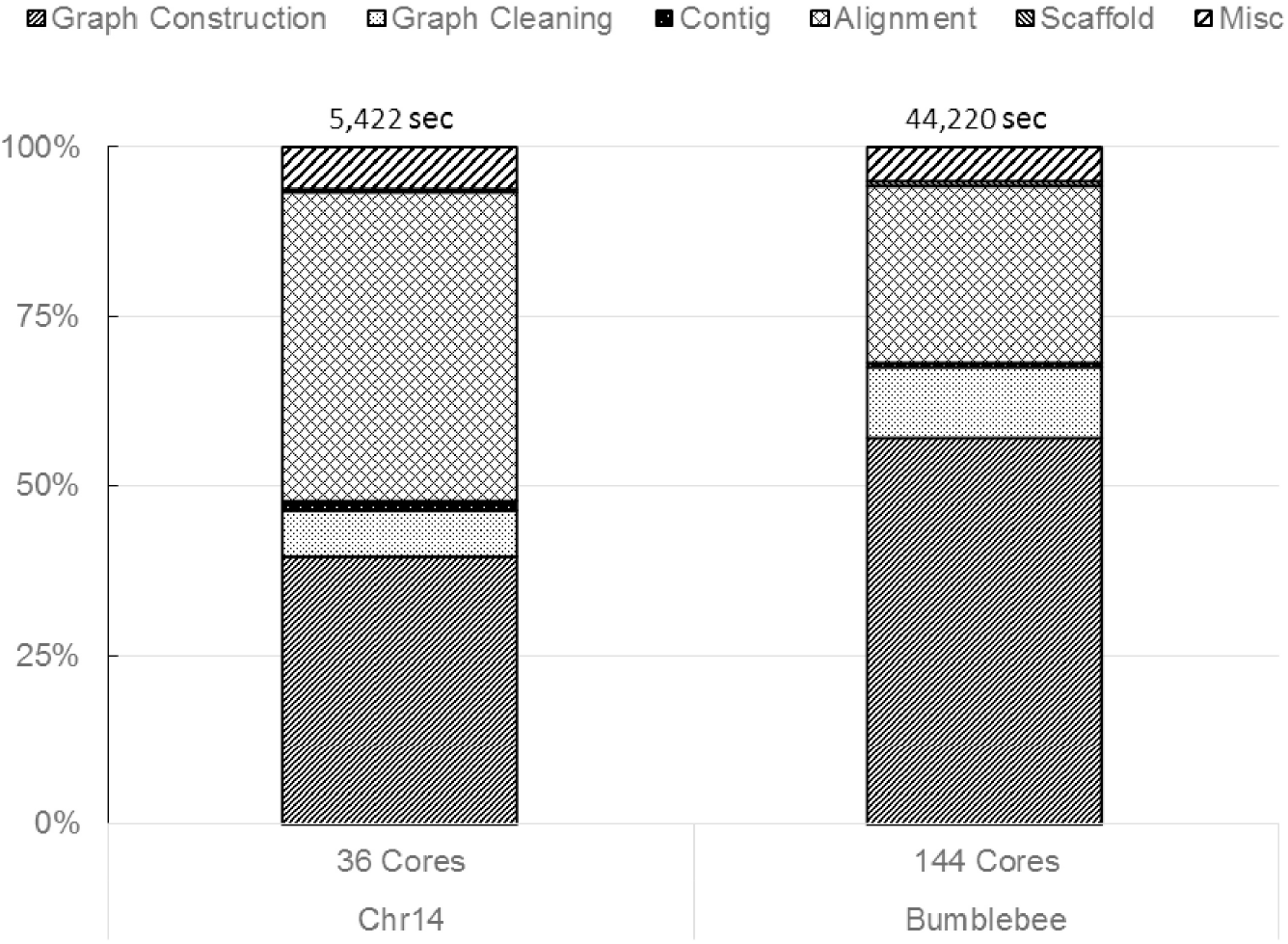
Percentage of overall runtime consumed by different blocks of ABySS on Chr14 and Bumble Bee datasets. The runtimes are collected by setting *k*-mer size to 51. The 144 core experiment uses 4 nodes of 36 cores each.

**Figure 5.**
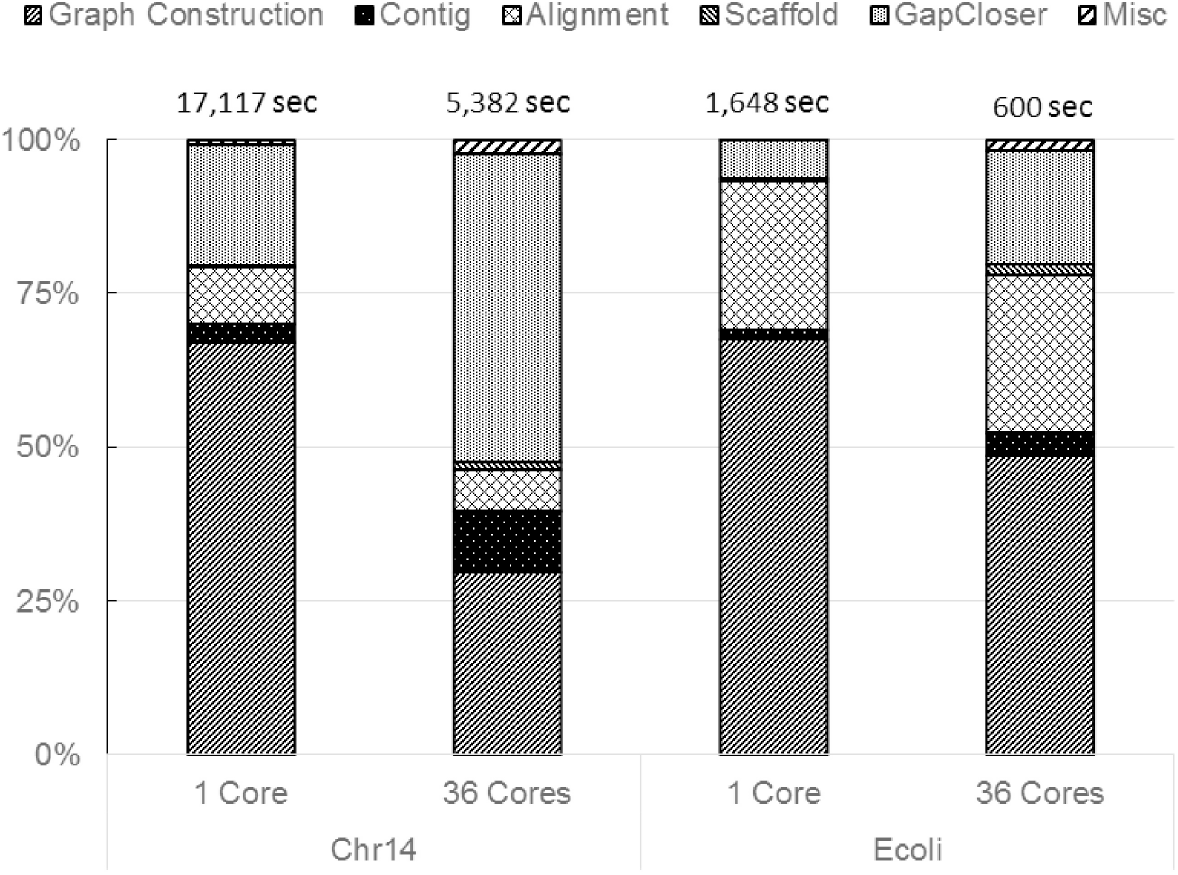
Percentage of overall runtime consumed by different blocks of SOAPDenovo2 on Chr14 and Ecoli datasets. The runtimes are collected by setting *k*-mer size to 51.

**Figure 6.**
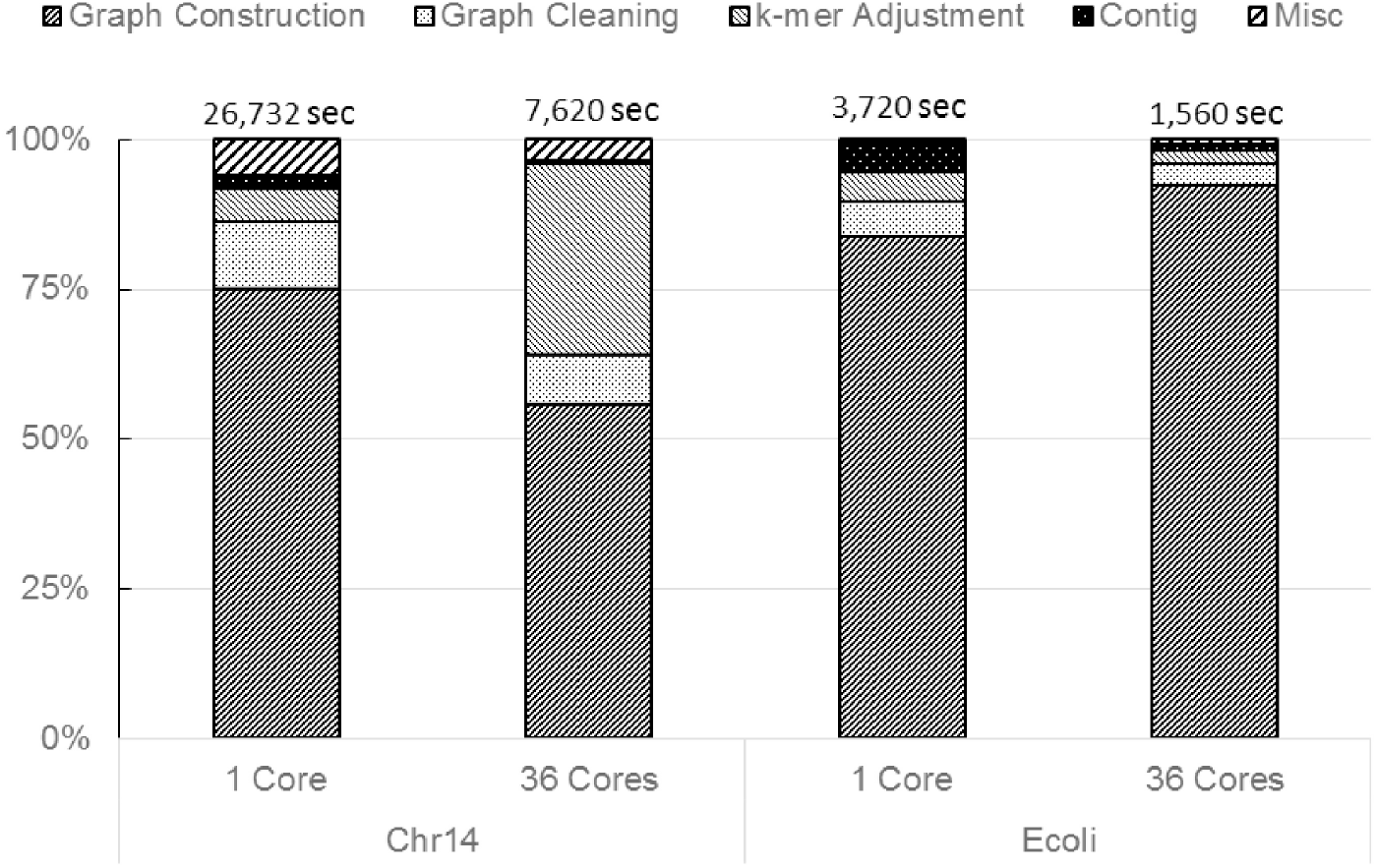
Percentage of overall runtime consumed by different blocks of SPAdes on Chr14 and Ecoli datasets.

**Figure 7.**
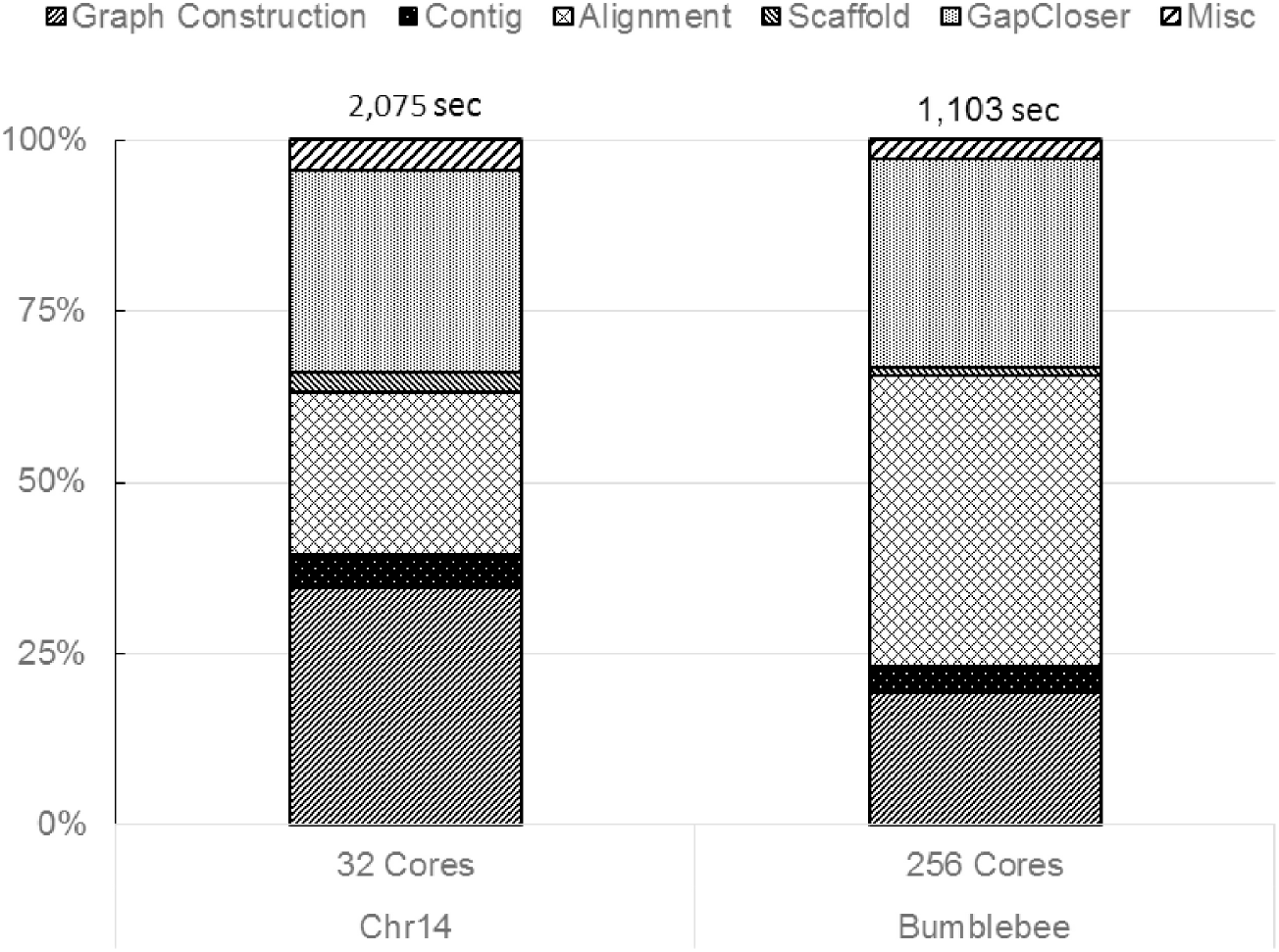
Percentage of overall runtime consumed by different blocks of HipMer on Chr14 and Bumble Bee datasets. The runtimes are collected by setting *k*-mer size to 51. The 256 core experiment uses 8 nodes of 32 cores each.

De Bruijn graph construction and sequence alignment are pervasive in genome assembly methods and consume the majority of the time. Excluding *gap closer*, these two blocks cover 63.9%-99.4% of the total time of these tools. Sequence Alignment is computationally similar to sequence mapping operation, thus, constituting of similar building blocks as the latter.

## 6 Variant Calling

Variant Calling is the process of identifying the differences between a sequenced genome and the corresponding reference. Variants can be in the form of single nucleotide (SNV), multiple nucleotides (MNV), insertion, deletion, or replacement. Among various available **VC** tools [12,17,41,55,57], **GATK**’s *HaplotypeCaller* (**HC**) for *germline* SNP/indel variant detection is among the most preferred. The popularity of **HC** is evident by the use of **GATK**’s *germline* workflow in various other **VC** workflows [15,23,28,51]. Hence, in this work, we exclusively focus on **HC** to find the building blocks of variant calling. Given a set of reads that are aligned to reference sequence, **HC** uses the following steps for **VC**.

### Active regions

**HC** narrows down the variant search space along the genome by finding the *active regions* which are potential regions in the genome likely to contain variants. *Active Regions* are identified as follows. First, a raw activity score is computed for each genome position, which realizes a raw activity profile. The raw activity score is the probability of a variant at a given position, calculated by reference-confidence model. Next, at each position *p*, all the activity scores over a small window of the genome with position *p* in the center are added to calculate the smoothed activity profile curve. Along the activity profile, local maxima that rise above a given threshold are located. Finally, appropriate intervals along the activity profile are set to extract the *active regions*.

### Re-assembly and haplotype extraction

Once the *active regions* are identified, the next goal is to construct the complete sample sequence (or haplotype) corresponding to each *active region*. These haplotypes are constructed by *de novo* assembly of all the reads that are mapped to the region, as follows. An assembly De Bruijn graph is constructed from the reference genome portion of the region. Then, all the reads corresponding to that region are passed along the paths in the assembly graph. For any mismatch, a new node is inserted into the assembly graph. Edges in the graph accumulate the support as the reads pass through them. Subsequently, haplotypes are extracted by traversing the paths that amassed enough support from the reads.

### Haplotype re-alignment

To identify the variant sites, for each active region, the haplotypes are re-aligned to the region in the reference sequence. This task is carried out using the Smith-Waterman algorithm.

### Haplotype evidence computation

The haplotype extraction step uses a quick heuristic based method to screen the haplotypes, and thus the extracted haplotypes act as candidate haplotypes to be verified later. Further evidence on haplotypes is gathered by aligning each read to the candidate haplotypes using Pairwise Hidden Markov Model (PairHMM) [19] algorithm. For a read and haplotype pair, PairHMM provides the likelihood score for the haplotype given the read. PairHMM also incorporates the base quality scores during likelihood calculations.

### Genotype assignment

**HC** performs the genotyping step, which classifies the variants in the haplotypes according to the genotypes. **HC** uses *Bayes theorem* to calculate the genotype likelihoods. Finally, **HC** reports all the identified variants in a **VCF** file.

#### 6.1 Results

We performed a detailed study of the source code of the **HC** tool from GATK-3.8 to understand the algorithm and identify boundaries of the above mentioned steps and hand instrumented it for profiling. **HC** was evaluated using the same settings used for evaluating the **GATK** best practices workflow. Figure 8 shows the assembly and SWA blocks to be runtime dominant. Note that the PairHMM step in **GATK-3.8** has already been accelerated using architecture-aware optimizations like SIMD based vectorization for modern multi-core processors. This is in contrast to the other key steps in the **NGS** tools studied in this work for which, while there have been significant efforts to improve the complexity of the algorithm, there is little architecture-aware programming to improve important performance determinants like data locality and number of instructions required. Thus, we also profiled **HC** with the unoptimized version of PairHMM so as to study it at the same level as the other key steps. Figure 8 shows runtime profiling results corresponding to optimized and unoptimized versions of the PairHMM step. Unoptimized PairHMM (UP) consumes ≈ 40% of the **HC** runtime. Thus, we also classify PairHMM as a key building block. The optimized version of PairHMM (OP) consumes only ≈ 11% of the **HC** runtime, thus improving the **HC** runtime by ≈ 18%. This result emphasizes that targeting significant building blocks for optimizations would result in correspondingly significant gains in the overall runtimes. *Misc* represents the un-profiled blocks and consumes a small portion of the overall runtime. Thus, the key blocks of assembly, PairHMM and SWA, cover 72% – 93% of the total runtime of variant calling.

**Figure 8.**
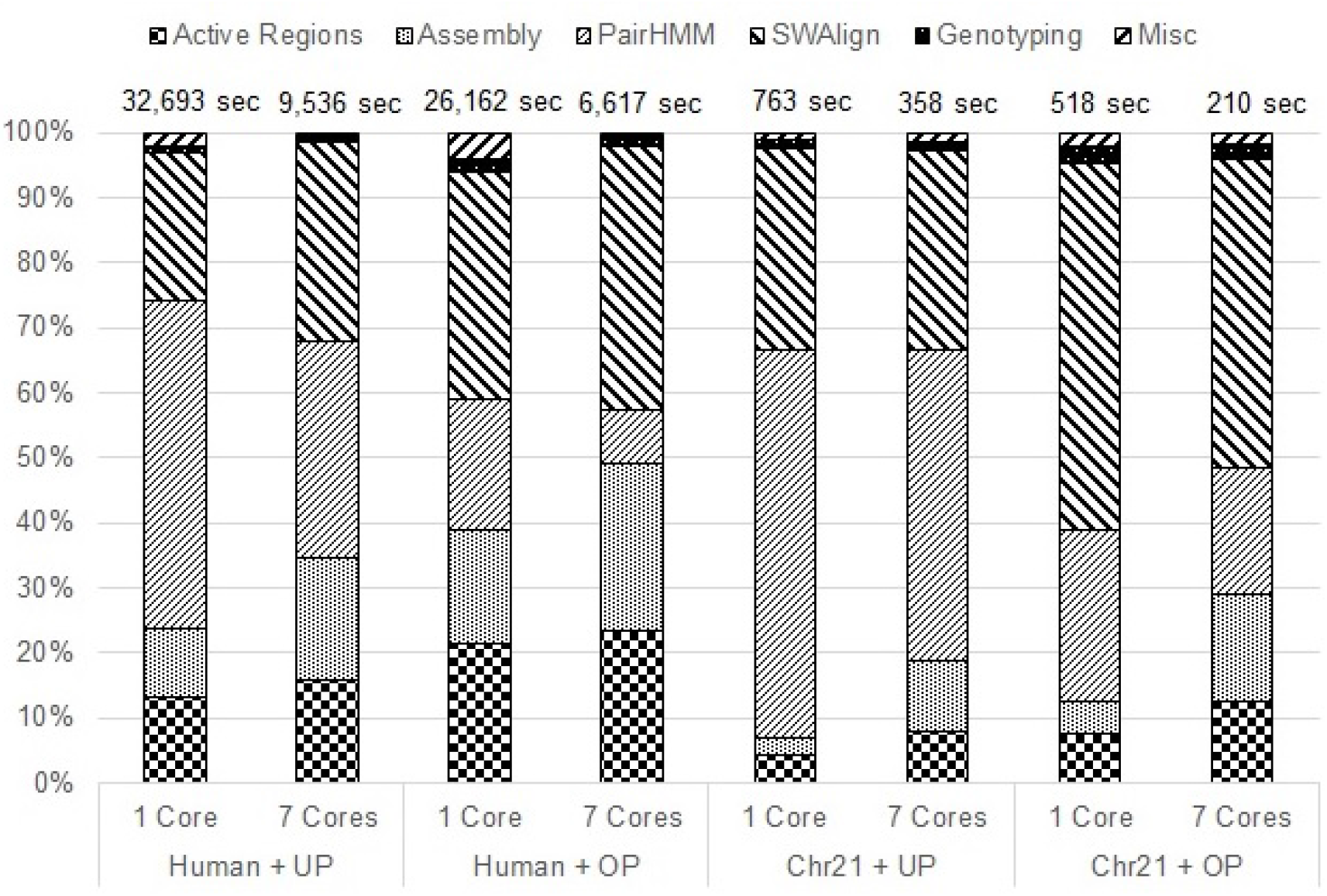
Percentage of the overall runtime consumed by different blocks of **HC**. Experiments are performed using optimized PairHMMM (OP) and un-optimized PairHMM (UP) on entire human genome and human chromosome 21 datasets.

## 7 Discussion

To summarize, we identified three important problems – sequence mapping, *De novo* assembly and variant calling – that account for a majority of the time consumed in **NGS** secondary analysis. For each of these, we studied the prominent tools in full detail understanding the source code and carving out key steps that are similar across tools.

We profiled these tools using real datasets to identify the most time consuming blocks. Our results show that sequence mapping spends a large portion of its time in **FM**-index based sequence search and Smith-Waterman algorithm. The most time consuming steps of *De novo* assembly are De Bruijn graph construction and sequence alignment. Sequence alignment, in turn, is very similar to sequence mapping and can be performed using one of the sequence mapping tools. Thus, it consists of the same building blocks as sequence mapping. A majority of time in variant calling is spent in assembly, Pairwise Hidden Markov Model algorithm, and Smith-Waterman algorithm. The assembly block of variant calling is performed using De Bruijn graphs and is similar in computation to *De novo* assembly. Apart from these computations, sorting is used quite frequently. It is one of the steps of **GATK** best practices workflow and is performed using Picard’s *SortSam* tool. It also appears frequently within quite a few tools.

Therefore, we identify four primary building blocks of **NGS** secondary analysis – **FM**-index based sequence search, Smith-Waterman algorithm, De Bruijn graph construction, and Pairwise Hidden Markov Model algorithm. We also identify sorting as a secondary building block. **FM**-index based sequence search exists in a few different flavors – exact and inexact match of seeds (default length 22) or entire reads and super maximal exact matches between the reads and the reference sequences. **BWA-MEM** uses banded Smith-Waterman algorithm without any need of backtracking information. On the other hand, HaplotypeCaller computes the full Smith-Waterman matrix and requires backtracking. De Bruijn graphs are constructed using single or multiple *k*-mer sizes and typically use hash tables for *k*-mer indexing and counting. PairHMM and SWA both use dynamic programming and are very similar in structure.

## 8 Conclusion and Future Directions

Given the rapid pace at which next generation sequencers are producing data, it is imperative to accelerate **NGS** secondary analysis. In this work, we performed a comprehensive study of secondary analysis methods. Our key finding is that the runtime is dominated by just four primary and one secondary building blocks. From our results, it is clear that any acceleration of these building blocks would go a long way in accelerating the overall execution of the **NGS** secondary analysis. Moreover, the fact that all the identified blocks are algorithmically mature puts us in a good position to do so. This work can help inform future hardware designs for the domain of Next Generation Sequencing. In addition, availability of standardized off-the-shelf implementation of such blocks that are optimized according to the hardware would not only accelerate current tools, but will also speed up the development of new tools in **NGS** secondary analysis.

## Key Points

- Availability of population genomic data has created opportunities for studying genomic differences across individuals or species.
- Advances in sequence data generation technology has prompted innovations in algorithmic and computational domain for the secondary analysis of sequence data.
- The progress in the NGS secondary analysis domain is marred both by the unavailability of suitable hardware and by incapability of the available tools in efficiently utilizing the available hardware.
- Majority of the secondary analysis tools make use of a few building blocks whose speedup can greatly increase the throughput of the tools or workflows using them.
- Hardware that is specifically tailored for these building blocks and implementations that can use that hardware optimally has the potential to improve the performance of NGS secondary analysis by leaps and bounds allowing it to keep pace with the rate of data generation.

## Acknowledgments

This research used resources of the National Energy Research Scientific Computing Center (NERSC), a U.S. Department of Energy Office of Science User Facility operated under Contract No. DE-AC02-05CH11231.

